# Sub-stoichiometric Degradation is Dispensable for Potent PROTACs: A Case Study for Irreversible Covalent BTK Degraders

**DOI:** 10.64898/2026.02.14.705950

**Authors:** Ran Cheng, Hanfeng Lin, Xin Yu, Shrilekha Misra, Andrew D. Mitchell, Jennifer A. Woyach, Xiaoli Qi, Jin Wang

## Abstract

Proteolysis-targeting chimeras (PROTACs) represent a transformative therapeutic modality, yet the viability of covalent PROTACs remains debated, as irreversible binding seemingly contradicts the catalytic mechanism central to their function. Here, we develop and characterize PSIRC3, a highly potent covalent PROTAC for Bruton’s tyrosine kinase (BTK) that addresses this ambiguity. PSIRC3 induces potent and selective BTK degradation with a sub-nanomolar DC_50_ of 0.75 nM and a D_max_ greater than 85%, while its non-covalent counterpart is completely inactive. This degradation activity is strictly dependent on covalent bond formation with the Cys481 residue, as evidenced by a total loss of efficacy against the C481S BTK mutant. PSIRC3 acts with remarkable speed, achieving maximum BTK degradation within 30 minutes, a kinetic profile linked to rapid cell permeation and efficient ternary complex formation. *In vivo*, a single administration of PSIRC3 leads to substantial BTK degradation in both PBMCs (>80%) and splenocytes (>50%). Computational modeling, parameterized with experimental data, reveals that degradation efficacy is governed by a delicate balance between E3 ligase and target protein affinities. Specifically, excessively high E3 affinity is detrimental by inducing a hook effect, while higher target affinity is generally beneficial. Our findings provide strong evidence that covalent engagement can drive potent and selective protein degradation, challenging the prevailing notion that catalytic turnover is indispensable for PROTAC efficacy. This work establishes a new benchmark for covalent degraders and opens new avenues for targeting previously intractable proteins.

## Introduction

Targeted protein degradation (TPD) has emerged as a revolutionary paradigm in drug discovery^1–3^. Central to this approach are proteolysis-targeting chimeras (PROTACs), bifunctional molecules that recruit a target protein to an E3 ubiquitin ligase, leading to ubiquitination and proteasomal degradation^4–6^. Unlike traditional inhibitors, PROTACs act through an event-driven mechanism: once a substrate is eliminated, the degrader is released to trigger additional rounds of degradation. This sub-stoichiometric mode of action is generally regarded as essential for developing potent degraders^7^.

This model, however, poses a mechanistic paradox for covalent PROTACs^8–10^. Covalent warheads are powerful tools for achieving strong and durable target engagement, particularly at shallow or cryptic binding sites^11–20^. While covalency on E3 ligases has demonstrated strong compatibility with the catalytic nature of targeted protein degraders^16,18,19,21–25^, the formation of irreversible bonds with target proteins is thought to impede catalytic turnover. As a result, each molecule may be limited to a single degradation event, thereby constraining overall potency^26–30^.

This debate has been clouded by ambiguous results from previous studies^6,30–33^. The most extensively studied target for comparing different binding modes is Bruton’s tyrosine kinase (BTK), the first kinase with an approved irreversible covalent inhibitor, ibrutinib. Ibrutinib binds to the kinase pocket of BTK and forms a covalent bond with C481 via a Michael addition reaction^34,35^. Consequently, a series of ibrutinib-based BTK PROTACs were developed and compared^31,36–39^. An early irreversible covalent BTK PROTAC from GSK failed to induce degradation, while its non-covalent counterpart was effective^36^. We also observed a similar phenomenon: the irreversible covalent BTK PROTAC IRC-1 displayed weaker degradation than the reversible non-covalent RNC-1 and the reversible covalent counterpart RC-1^37^. Both studies suggested that irreversible PROTACs are not potent, possibly due to a lack of catalytic function. Irreversible BTK PROTACs IR-2 developed by the London group and another by the Calabrese group at Pfizer were potent but complicated because they were active against both wild-type (WT) and C481S mutant BTK^38,39^. The fundamental limitation of these studies was their reliance on the ibrutinib warhead, which possesses high non-covalent binding affinity that masks the true contribution of the covalent bond formation. This has made it difficult to deconvolute the specific role of covalency in PROTAC-mediated degradation. Similarly, although engineered systems such as HaloTag-targeting chloroalkane degraders (HaloPROTACs) have proven valuable for investigating target functionality^40,41^, they often involve overexpression of the E3 ligase, which may artificially enhance the observed potency. The E3 ligase—particularly CRBN—frequently constitutes the limiting factor in concentration between the target and E3, depending on the specific system under study. All these issues made it difficult to deconvolute the specific role of covalency in PROTAC-mediated degradation in a therapeutic relevant setting.

To address this challenge, we leveraged poseltinib^42^, a covalent BTK inhibitor whose binding is strictly dependent on covalent bond formation between BTK Cys481 and the warhead. Unlike ibrutinib, poseltinib shows no appreciable affinity for the C481S mutant, and its non-covalent analog fails to bind WT BTK (**Figure 1**), providing a clean system to study covalent PROTACs without interference from non-covalent interactions. Here, we report PSIRC3, a poseltinib-based covalent PROTAC that achieves highly potent, selective, and rapid degradation of BTK. Computational modeling of PROTAC kinetics revealed that degradation potency is governed by a delicate balance between E3 ligase and target protein affinities. Our simulations show that excessively high E3 affinity is detrimental, inducing a “hook effect,” while higher target affinity is generally beneficial, with overall efficacy being modulated by cellular protein concentrations. Our findings demonstrate that covalent engagement can serve as the sole driver of efficient targeted protein degradation, definitively establishing covalent PROTACs as a viable and broadly applicable degrader modality.

**Figure 1.**
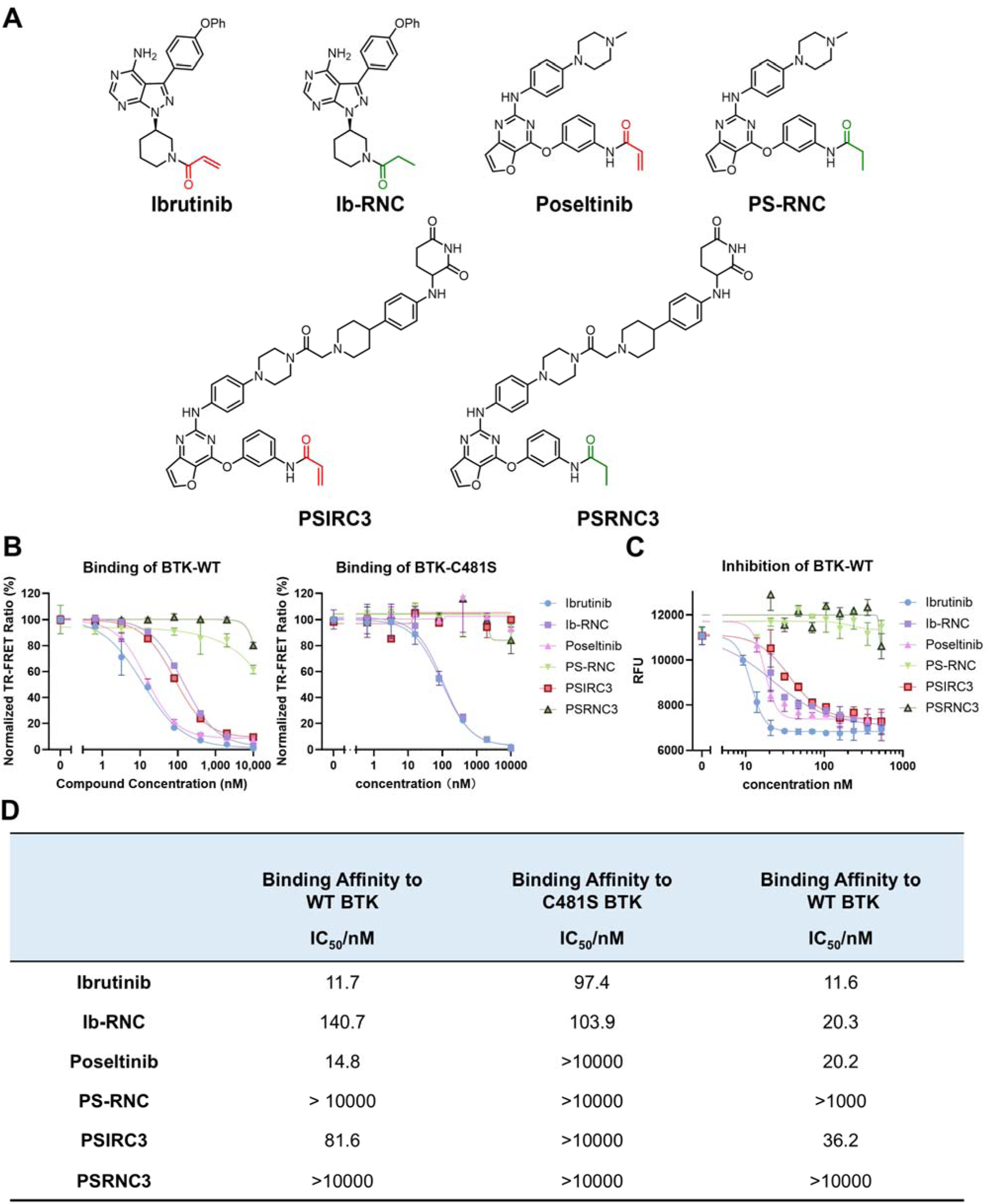
Characterization of Covalent BTK PROTACs. (A) Chemical structures of covalent BTK inhibitors (ibrutinib, poseltinib), their non-covalent counterparts (Ib-RNC, PS-RNC), the covalent PROTAC PSIRC3, and its non-covalent control PSRNC3. (B). TR-FRET based binding assay between compounds and WT-BTK and C481S BTK protein. Serial dilutions of compounds mixed with 2 nM of His-BTK, 0.3 nM Tb-anti-His, and 120 nM of BTK-BODIPY tracer. (C) The biochemical WT-BTK inhibition with 1h inhibitor inhibition (BTK Inhibition IC50) was measured using the kinase assay kit from AssayQuant Technologies Inc. (D) Binding affinity data from TR-FRET assay and inhibition data from kinase activity assay.

## Results and Discussion

### Validation of a Covalency-Dependent Warhead for BTK

The success of ibrutinib, the first FDA-approved covalent inhibitor of Bruton’s tyrosine kinase (BTK), paved the way for exploring covalent PROTACs. However, ibrutinib’s utility as a mechanistic tool is limited. Despite its covalent bond with Cys481 (**Figure 1A**), it retains significant non-covalent binding affinity, showing only an ∼10-fold decrease in binding to the C481S mutant (**Figure 1B-C**). Its Michael acceptor-saturated analog, Ib-RNC (Reversible Non-Covalent), also remains binding to wild-type BTK (**Figure 1B-C**). This non-covalent activity, also seen in kinase assays (**Figure 1C, Figure S1**), implies that ibrutinib-based covalent PROTACs may have confounding non-covalent characteristics.

To isolate the role of covalency, we turned to poseltinib. In stark contrast to ibrutinib, poseltinib’s binding is strictly covalent-dependent. It displayed robust binding to WT BTK but completely lost affinity for the C481S mutant (**Figure 1B-C**). Furthermore, its saturated analog, PS-RNC, showed no detectable binding to WT BTK. This covalent dependency was also mirrored in kinase inhibition assays (**Figure S1**), where poseltinib potently inhibited WT-BTK, while PS-RNC was inactive. Both poseltinib and PS-RNC were inactive against the C481S mutant BTK. This confirms poseltinib as an ideal, “clean” warhead for covalent BTK PROTACs, whose engagement is exclusively driven by covalent bond formation.

### Design and Biochemical Characterization of Covalent PROTAC PSIRC3

Leveraging this covalency-isolated warhead, we designed the covalent PROTAC PSIRC3 (IRreversible Covalent) and its non-covalent analog PSRNC3 as a negative control (**Figure 1A**). We next evaluated whether PSIRC3 biochemically inherits the strict covalent dependency of its poseltinib warhead.

Indeed, the binding and inhibition profiles mirrored those of the warhead. In TR-FRET assays^37,43,44^, PSIRC3 exhibited potent binding to WT-BTK with an IC of 81 nM (**Figure 1B, 1D**). In contrast, its binding was completely abolished against the C481S mutant (**Figure 1B, 1D**). The non-covalent control, PSRNC3, which lacks the electrophilic acrylamide, showed no binding to either WT or C481S BTK (**Figure 1B, 1D**).

Kinase inhibition assays confirmed this covalent requirement. PSIRC3 potently inhibited WT-BTK, while its non-covalent control PSRNC3 was inactive (**Figure 1C, Figure S1**). As expected, both PSIRC3 and PSRNC3 were inactive against the C481S mutant (**Figure 1C, Figure S1**).

To provide definitive proof of the covalent mechanism, we also performed intact protein mass spectrometry. Incubation of purified BTK protein with PSIRC3 resulted in a mass shift corresponding precisely to the adduction of the PROTAC molecule (Δ=783 Da), confirming covalent bond formation. No such modification was observed with the non-covalent counterpart, PSRNC3 (**Figure 2**). These data demonstrate that PSIRC3 is a *bona fide* covalent-dependent PROTAC at the biochemical level, with target engagement being strictly reliant on its ability to modify BTK C481.

**Figure 2.**
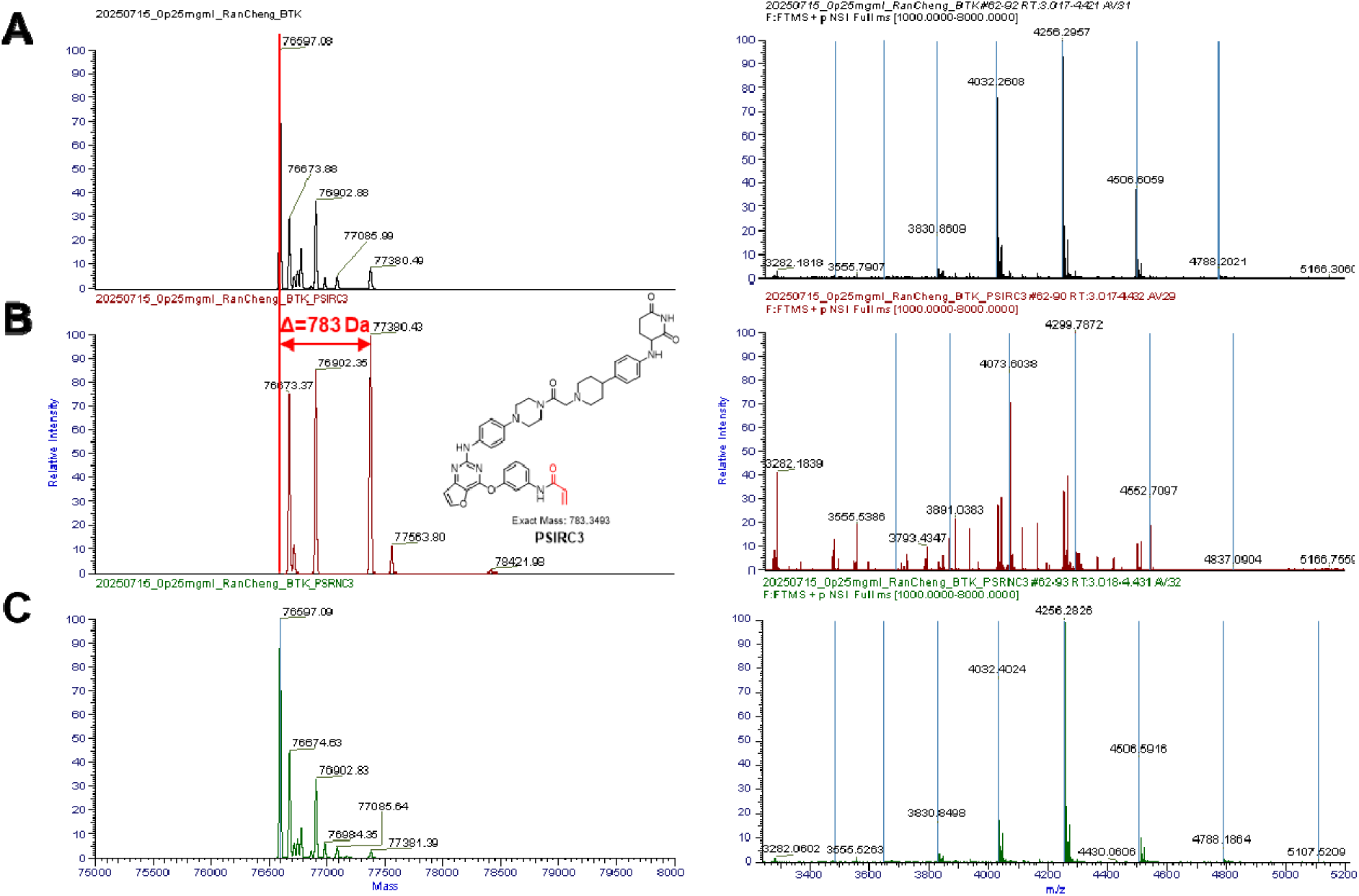
Direct Confirmation of Covalent Target Modification by Mass Spectrometry. Deconvoluted mass spectra (left) and raw mass spectra (right) of purified BTK protein after incubation with (A) DMSO vehicle, (B) PSIRC3, and (C) PSRNC3. A distinct mass shift of 783 Da, corresponding to the molecular weight of PSIRC3, is observed exclusively in panel B, confirming the formatio covalent adduct.

### Covalent Engagement by PSIRC3 Drives Potent Cellular Degradation

Having established PSIRC3 as a covalent-dependent binder, we next assessed its ability to induce protein degradation in cells. We conducted comprehensive degradation assays using BTK-HiBiT Ramos and BTK-nLuc Ramos cell lines, utilizing bioluminescence-based methods to quantify BTK levels. PSIRC3 demonstrated exceptional potency in BTK-HiBiT Ramos cells, exhibiting a sub-nanomolar DC_50_ of 0.75 nM and achieving a remarkable 85% D_max_ in BTK-HiBiT Ramos cells (**Figure 3A**). We confirmed the degradation mechanism occurs via the ubiquitin-proteasome pathway (**Figure 3D**). Similarly, robust degradation activity was observed in BTK-nLuc Ramos cells, with a DC_50_ of 1.5 nM. In stark contrast, the non-covalent control PSRNC3 displayed negligible degradation potency in both cell lines (**Figure 3A**).

Western blot analysis corroborated these results. Ramos cells treated with PSIRC3 and PSRNC3 for 24 hours showed potent, dose-dependent BTK degradation by PSIRC3 (DC_50_ of 5.6 nM), while PSRNC3 failed to induce any appreciable degradation (**Figure 3B**).

**Figure 3.**
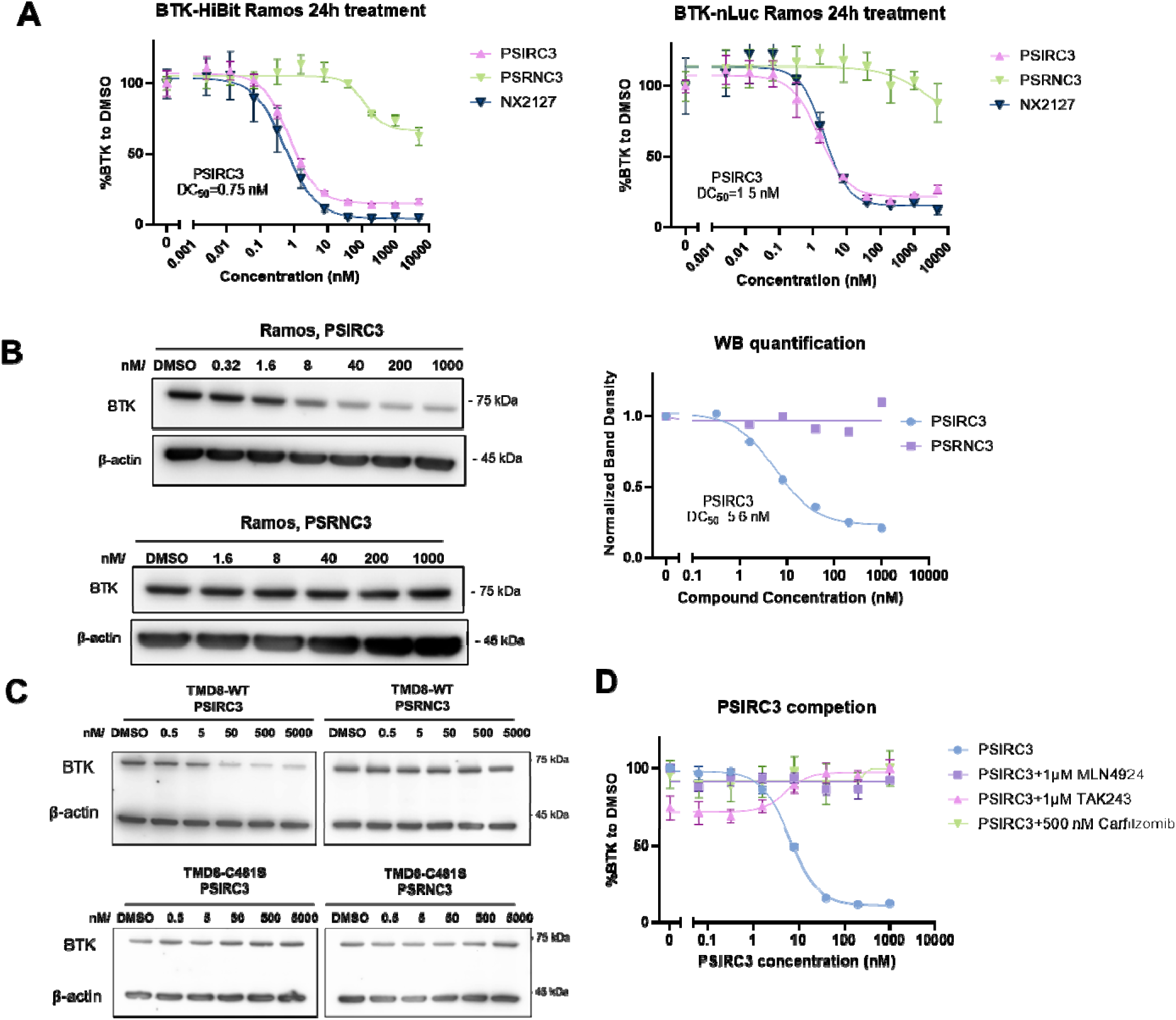
PSIRC3-induced BTK degradation is Dependent on the Covalent Bond Formation. **(A)** BTK-HiBiT Ramos cells and BTK-nLuc Ramos cells were treated with indicated compounds for 24 h. The BTK degradation was determined by evaluating luminescence signals of NanoLuc. Shown are mean ± SD luminescence values normalized to DMSO-treated samples. **(B)** Ramos cells were treated with indicated compounds at 0, 0.32, 1.6, 8, 40, 200, and 1000 nM for 24 h, followed by Western blotting for BTK. **(C)** TMD8-WT and TMD-C481S cells were treated with indicated compounds at 0, 0.5, 5, 50, 500, and 5000 nM for 24 h, followed by Western blotting for BTK. **(D)** Mechanistic validation of degradation in BTK-HiBiT Ramos cells. Cells were pre-treated for 1h with the E1 inhibitor TAK243, NEDDylation inhibitor MLN4924, or proteasome inhibitor carfilzomib prior to PSIRC3 treatment. Data are normalized to DMSO control and represent mean ± SD.

To further confirm that this cellular activity was driven by the covalent mechanism, we used wild-type and C481S mutant BTK-expressing cell lines. Parental TMD8 cells with wild type BTK and TMD8 cells with BTK-C481S mutant engineered via CRISPR-Cas9 technology were treated with PSIRC3 and PSRNC3 for 24 hours. Western blot analysis (**Figure 3C**) showed that PSIRC3 induced potent BTK degradation in wild-type TMD8 cells but had no degradation potency in C481S-BTK TMD8 cells. PSRNC3 exhibited no detectable protein degradation in either cell line.

The results provide clear evidence that covalent bond formation at BTK Cys481 is essential for PSIRC3 to induce effective cellular BTK degradation. This supports the classification of PSIRC3 as a genuine irreversible covalent PROTAC and challenges assumptions regarding the inferiority of covalent PROTACs due to non-catalytic degradation, thereby underscoring their promise as a significant class of therapeutic agents.

### Irreversible Binding Creates a Kinetic Trap that Outcompetes Reversible Inhibitors

To demonstrate the functional consequence of this irreversible binding in a competitive cellular environment, we performed a target competition assay. BTK-HiBiT Ramos Cells were pre-treated for 12 hours with 100 nM of the covalent BTK inhibitor ibrutinib or the non-covalent BTK inhibitors ARQ531 or LOXO305 to occupy the target’s binding site. Then, varying concentrations of either the covalent PROTAC PSIRC3 (**Figure 4A**) or the non-covalent PROTAC NX-2127^45^ (**Figure 4B**) were added in the continued presence of the inhibitors for up to 48 hours.

**Figure 4.**
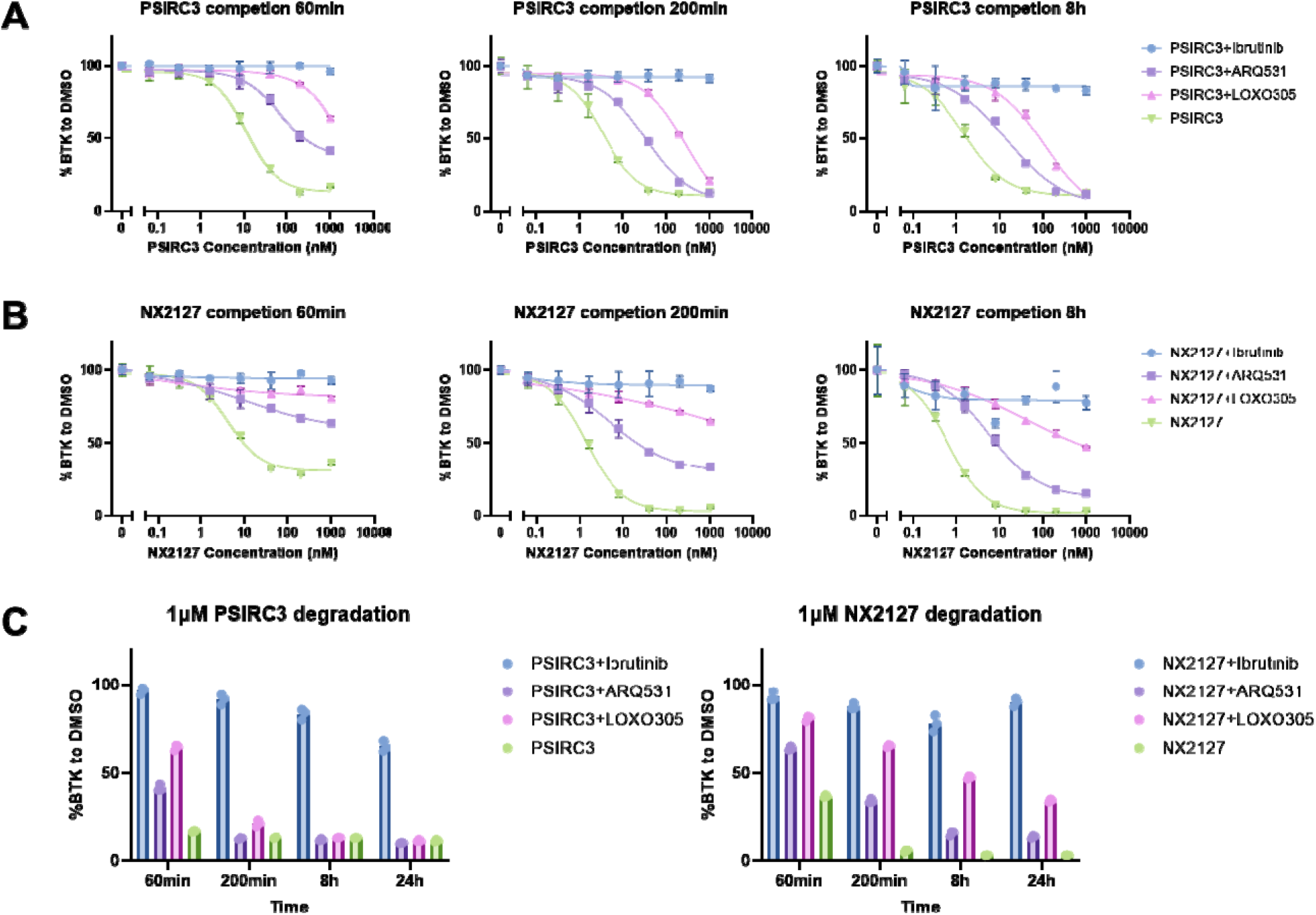
Functional Validation of Irreversible Action via Target Competition Assay. Quantification of BTK degradation over time in cells pre-treated with covalent and non-covalent BTK inhibitor for 12 h, followed by the addition of either **(A)** PSIRC3 (covalent) or **(B)** a non-covalen BTK PROTAC NX-2127. **(C)** Comparison of BTK degradation induced by PSIRC3 or NX-2127 over time (60 min to 24 h) in the presence of competitive inhibitors.

As shown in **Figure 4B**, the degradation activity of the non-covalent PROTAC, NX-2127, was significantly hindered by all three pre-treated inhibitors, with the dose-response curve shifting substantially to the right. In contrast, our covalent PROTAC, PSIRC3, demonstrated a superior ability to overcome competition from the *non-covalent* inhibitors (ARQ531, LOXO305) (**Figure 4A**). The dose-response curves for PSIRC3 showed a much smaller rightward shift. As ibrutinib irreversibly occupies the Cys481 residue, neither PSIRC3 nor NX-2127 could induce BTK degradation, serving as an important control.

To further highlight the kinetic advantage of PSIRC3, we analyzed degradation efficacy at a single 1 μM concentration over 24 hours (**Figure 4C**). When competing with the non-covalent inhibitors (ARQ531 and LOXO305), PSIRC3 achieved near-maximal degradation within 8 hours. This rapid action demonstrates its ability to quickly and permanently occupy the BTK binding site via covalent bond formation. Conversely, NX-2127 showed very limited degradation even after 24 hours, confirming its inability to effectively compete with the pre-bound non-covalent inhibitors. These findings indicate that the irreversible binding of PSIRC3 enables it to continuously target and degrade BTK as reversible inhibitors gradually dissociate, ultimately resulting in complete protein degradation. This highlights a crucial functional advantage of the covalent mechanism. As a result, covalent PROTACs like PSIRC3 may offer greater durability and achieve deeper target knockdown, making them particularly promising for use as combination therapies in patients who are already receiving reversible inhibitors but have not achieved a full response. In contrast, reversible PROTACs are unable to be combined effectively with BTK inhibitors.

### Rapid degradation by PSIRC3 is driven by efficient ternary complex formation

To understand the kinetic drivers of degradation, we compared PSIRC3 to two structural analogs, PSIRC6 and PSIRC8. All three compounds share the same covalent warhead and E3 ligase binder, differing only in their linker (**Figure 5A**). Despite this, their degradation activities were strikingly different. PSIRC3 induced potent and rapid degradation, achieving D_max_ within 30-60 minutes (**Figure 5B**). In contrast, PSIRC6 and PSIRC8 are much weaker and slower degraders, requiring over 4 hours to reach maximal effects.

**Figure 5.**
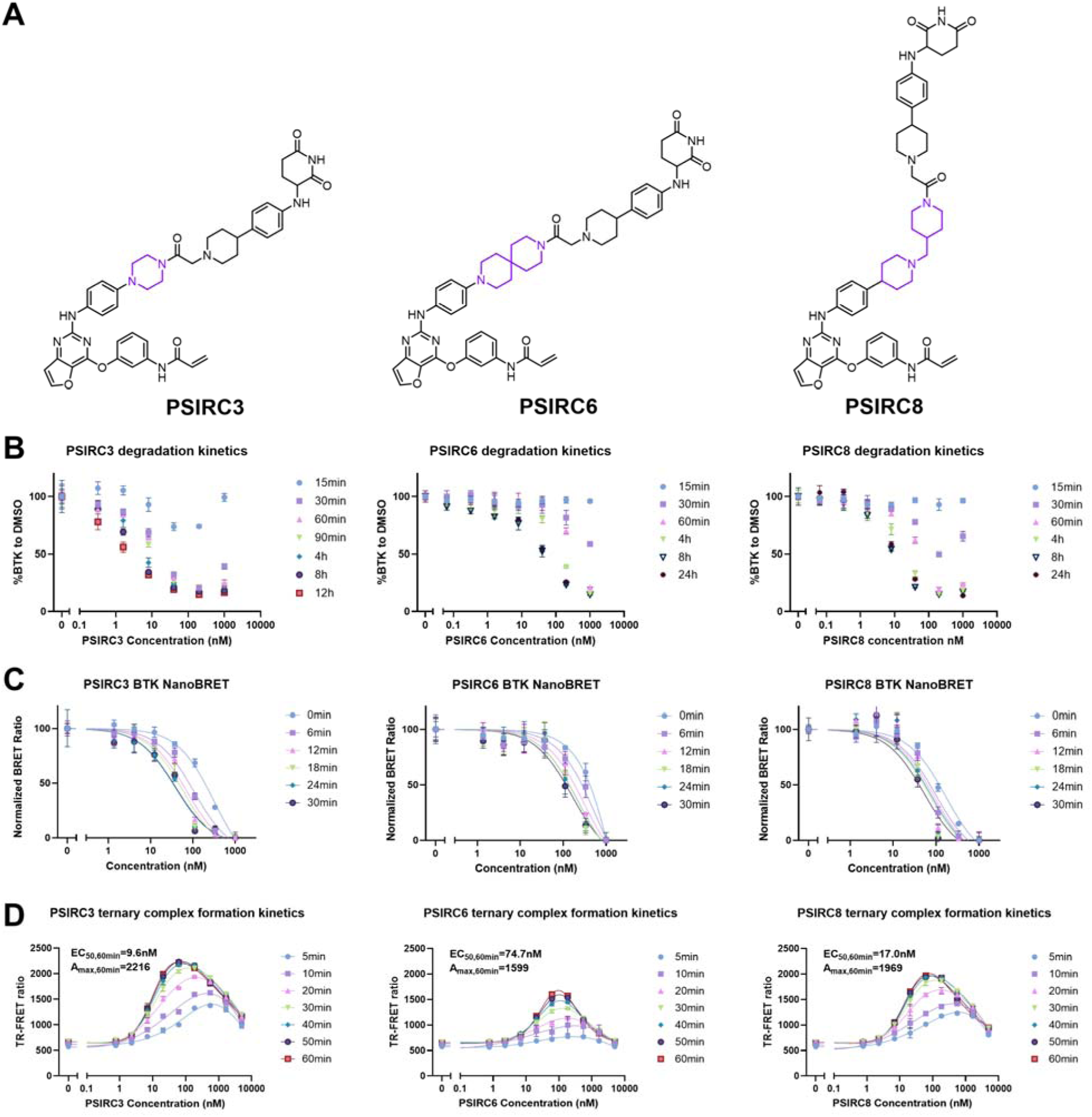
Correlation Between BTK Degradation Kinetics, in-cell target engagement. **(A)** Chemical structures of PSIRC3, PSIRC6, and PSIRC8 with the same warhead and E3 ligand, but different linkers. **(B)** BTK-HiBiT Ramos cells were treated with PSIRC3, PSIRC6, PSIRC8 for indicated time, the BTK degradation was determined by evaluating luminescence signals of NanoLuc. **(C)** BTK in-cell target engagement assay. BTK-nLuc Ramos cells treated with a BTK NanoBRET tracer, which binds to BTK-nLuc to induce bioluminescence resonance energy transfer (BRET). Adding PSIRC3/6/8 to cells would compete with the CRBN tracer binding to CRBN, thus reducing the NanoBRET signals. **(D)** Formation of BTK-PROTAC-CRBN ternary complex induced by PSIRC3/6/8. PROTACs bring Tb antibody-labeled BTK and fluorescence-labeled His-CRBN/DDB1 into proximity and results in TR-FRET signal increase.

We hypothesized this difference could be due to variations in cell permeability, target engagement, or ternary complex formation. First, we tested cell entry and target engagement using a NanoBRET assay. This showed that all three compounds—PSIRC3, PSIRC6, and PSIRC8—rapidly entered the cell and engaged BTK with nearly identical kinetics, reaching equilibrium within 20 minutes (**Figure 5C**). This result demonstrates that cellular permeability and target binding are not the major distinguishing factors.

The critical difference was revealed in the ternary complex formation assay (**Figure 5D**). The compounds’ ability to form a BTK-PROTAC-CRBN complex correlated strongly with their degradation potency. PSIRC3 induced a potent, dose-dependent, and rapid ternary complex signal, with a EC_50_ of 9.6 nM and Amax of 2216 at 60 min. PSIRC8 and PSIRC6 induced the formation of fewer ternary complexes, with a EC_50_ of 74.7 nM and 17nM, respectively. This powerful comparative data (Degradation: PSIRC3 > PSIRC8 > PSIRC6; Ternary Complex: PSIRC3 > PSIRC8 > PSIRC6) supports that the linker’s ability to drive a cooperative and productive ternary complex, not just cell entry or target binding, is the key determinant of potent covalent degradation.

### PSIRC3 Exhibits Potent *in vivo* Efficacy and High Selectivity

We observed that PSIRC3 exhibits significant *in vivo* efficacy in mice. Following a single intravenous (I.V.) injection of PSIRC3 at 15 mg/kg, peripheral blood mononuclear cells (PBMCs) and splenocytes were harvested 4 hours post-injection. BTK levels in these cells were quantified via western blot (**Figure 6A**). PSIRC3 induced substantial BTK degradation, with greater than 80% reduction in PBMCs and over 50% reduction in splenocytes. Although demonstrating *in vivo* efficacy is not the main focus of this work, this proof-of-concept experiment supports the potential of PSIRC3 as a covalent PROTAC and highlights the promise of this platform for achieving potent protein degradation *in vivo*.

**Figure 6.**
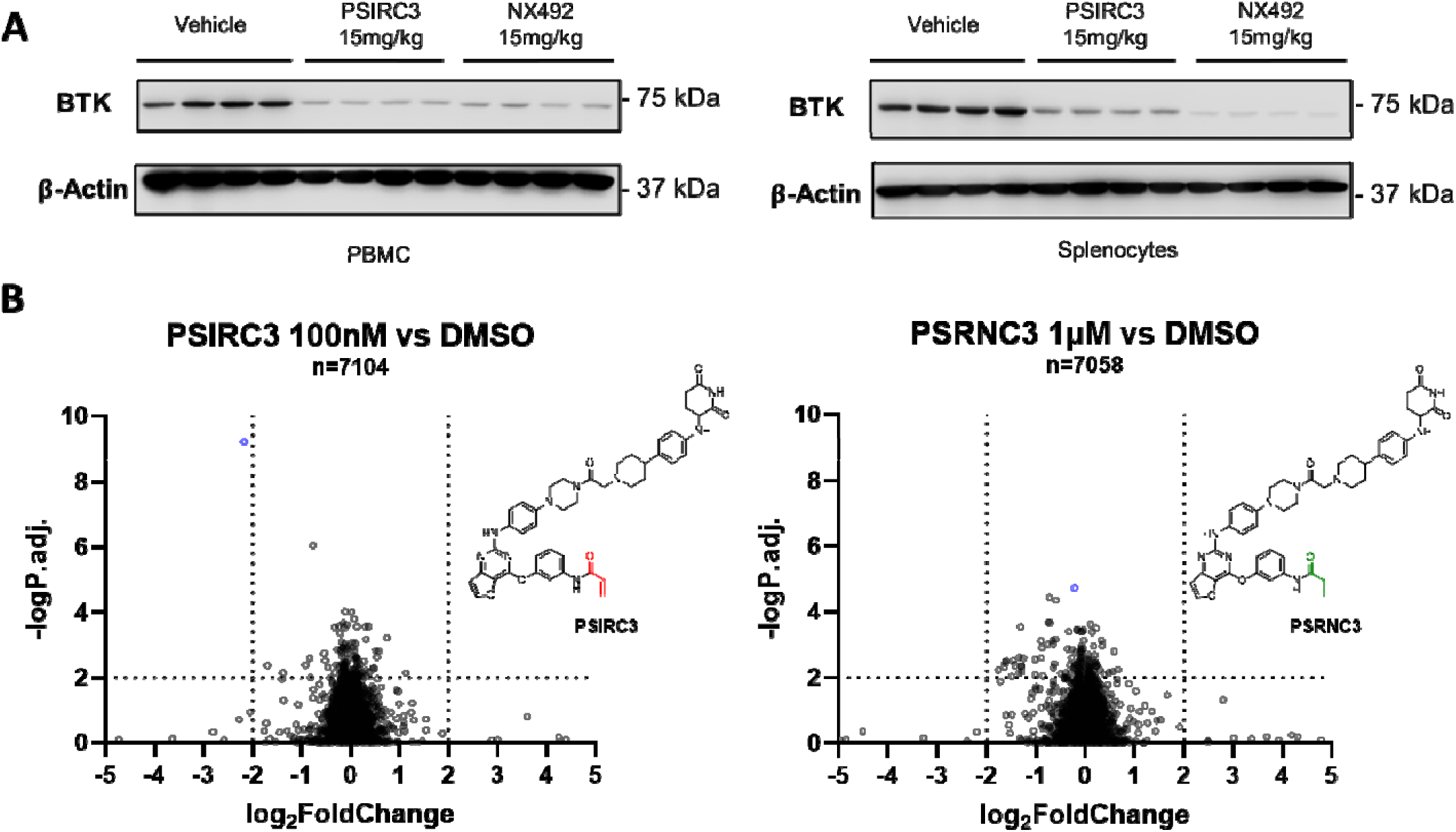
PSIRC3 Shows BTK Degradation *in vivo* and High Selectivity in Proteomics. **(A)** PBMCs and spleens were extracted from BALB/c mice after a 4h I.V. injection and immunoblotted for BTK. **(B)** Volcano plots from quantitative mass spectrometry-based proteomics of Ramos c lls treated with PSIRC3, PSRNC3, or DMSO. The plot shows log2 fold change versus -log10 adjusted p-value for all quantified proteins, highlighting BTK as the most significantly downregulated target.

Consistent with the observed *in vivo* efficacy, we further evaluated the degradative potential of PSIRC3 in primary cells derived from patients. In B cells isolated from chronic lymphocytic leukemia (CLL) patients (n=4), PSIRC3 treatment induced a dose-dependent degradation of BTK (**Figure S2A**). Substantial protein reduction was observed at concentrations as low as 5 nM, with near-complete depletion achieved at higher doses across all patient samples. Consistent with BTK degradation, we observed a downregulation of key downstream effectors governing cell survival (*BCL-XL*), proliferation (*MYC*, *OCT2*), B-cell activation (*CD40*), and microenvironmental homing (*CXCR4*, *CXCR5*, and *CXCR7*) (**Figure S2B**). This activity in clinical specimens underscores the translational potential of PSIRC3 in treating BTK-dependent malignancies.

To evaluate selectivity, we conducted a global proteomic analysis in Ramos cells treated with PSIRC3 and PSRNC3 for 12 hours. This experiment quantified changes in the levels of over 7,000 proteins. As shown in the volcano plots (**Figure 6B**), our analysis revealed that PSIRC3 selectively degraded BTK with high specificity. This high degree of selectivity is a crucial attribute for a therapeutic agent. Meanwhile, PSRNC3 showed little BTK degradation, as expected.

### Lessons from covalent PROTAC design

To systematically dissect the complex kinetic parameters governing covalent PROTAC efficacy, we developed a computational model using MATLAB SimBio software. The model was parameterized using initial conditions derived from our experimental data, including the cellular concentrations of the target protein (T) and E3 ligase (L) (**Figure S5**), while varying PROTAC (P) concentration. This model simulates key events including binary and ternary complex formation, ubiquitination, and degradation (**Figure 7A-B**). By varying key parameters, we gained critical insights into the distinct behaviors of covalent versus non-covalent degraders.

**Figure 7.**
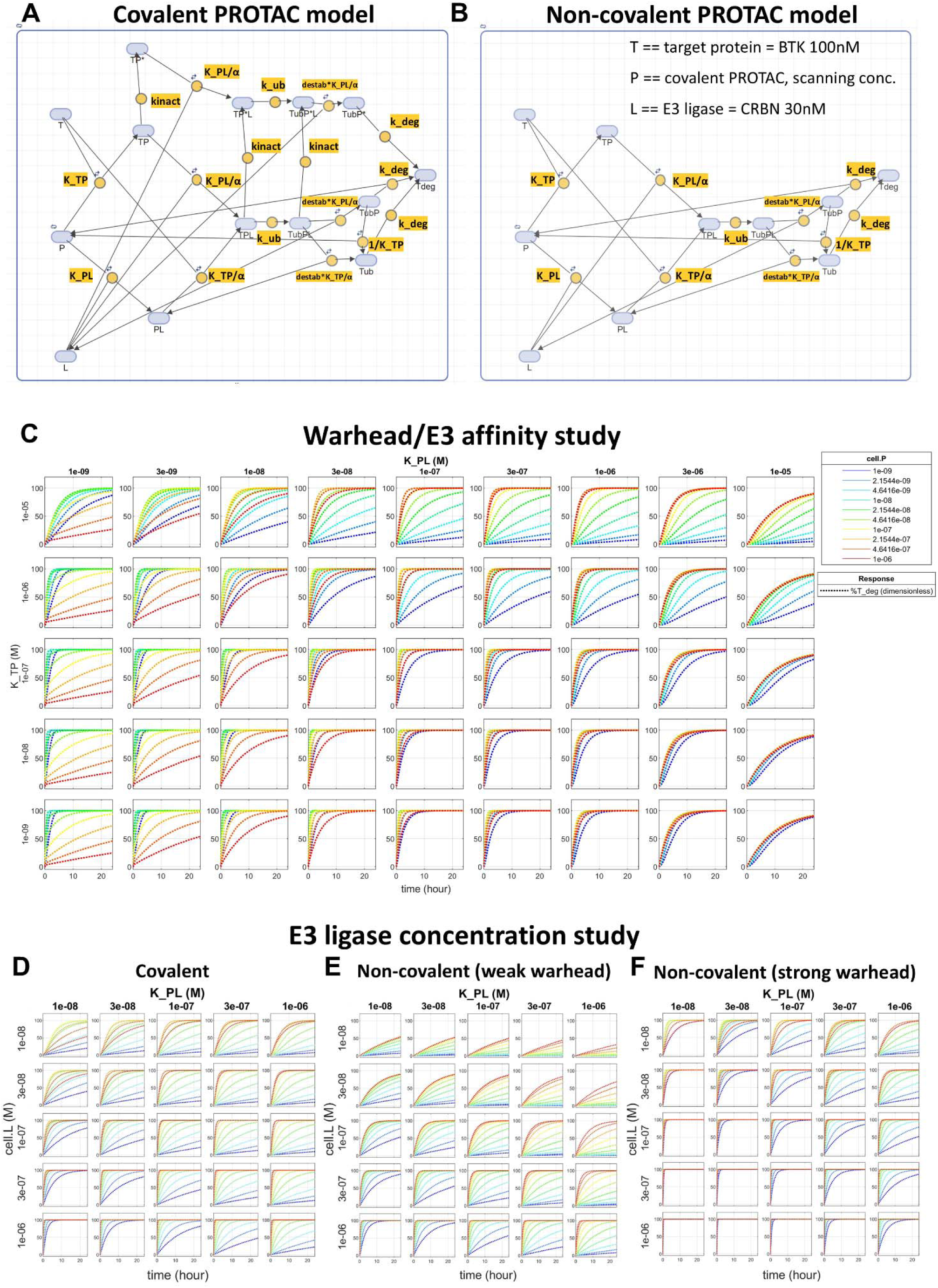
SimBio Modeling of Both Covalent and Non-covalent PROTAC Degradation Events. **(A-B)** covalent and non-covalent PROTAC mechanism models in which target protein (T) undergoes ternary complex (TPL) formation with PROTAC (P) and E3 ligase (L), followed by ubiquitination (Tub) and degradation (Tdeg). Asterisk denotes the presence of covalent bond between T and P (e.g. TP*, TP*L). All the parameters (kinetic constant if an irreversible step, equilibrium constant if a reversible step) with their expressions are highlighted in yellow. **(C)** The 24 hrs target protein degradation kinetics simulations for covalent PROTAC (1 nM to 1 μM) with varying warhead affinity (K_TP) and E3 ligase affinity (K_PL), and all other kinetic parameters as constants. Results are presented as percentage of degraded target protein (%T_deg). **(D-F)** The 24 hrs target protein degradation kinetics simulations for both covalent PROTAC (D: warhead K_TP = 1e-5 M, kinact = 0.01 s^-1^) and non-covalent PROTAC (E: weak warhead K_TP=1e-5 M; F: strong warhead K_TP=1e-7 M) (1 nM to 1 μM) with varying E3 ligase affinity (K_PL) and E3 ligase concentrations (cell.L). Results are presented as percentage of degraded target protein (%T_deg).

#### The Critical Balance of E3 Ligase and Target Affinity

A central challenge in PROTAC design is the hook effect. Our simulations for covalent PROTACs revealed that E3 ligase affinity (K_PL) is a primary determinant of this phenomenon. Contrary to the intuition that tighter binding is always better, our model shows that excessively high affinity for the E3 ligase is detrimental, leading to a severe hook effect (**Figure 7C**). This is attributed to the high-affinity binding sequestering the ligase into unproductive PROTAC-E3 binary complexes. This depletion of free E3 ligase stalls the degradation pathway, leading to the accumulation of the covalent Target-PROTAC binary adduct (TP*) (**Figure S3C**). This modeling insight aligns with our experimental observation that the hook effect for PSIRC3 attenuated over time (**Figure 5B**), suggesting it is a kinetic bottleneck. The E3 ligase affinity scanning suggests that the CRBN binder used in PSIRC3 fortuitously occupies an affinity ‘sweet spot’—potent enough for efficient ternary complex formation but not so high as to induce a debilitating hook effect.

In contrast to E3 ligase affinity, the model indicates that a higher target protein binding affinity (K_TP) is almost always beneficial, promoting faster and more complete degradation. However, even a very strong warhead cannot overcome the potent hook effect induced by an overly tight E3 binder, underscoring the delicate balance required (**Figure 7C**).

#### Contrasting Covalent and Non-Covalent PROTAC Models

The simulations also highlighted key differences between covalent and non-covalent PROTACs. For non-covalent PROTACs, the model predicts that a weaker warhead can paradoxically tolerate a much more potent E3 ligase binder without inducing a significant hook effect (**Figure 7E**). Conversely, a non-covalent PROTAC with a strong warhead behaves similarly to a covalent PROTAC, exhibiting a pronounced hook effect when paired with a high-affinity E3 binder (**Figure 7F**). This convergence occurs because a very high-affinity, slow-dissociating non-covalent interaction begins to kinetically approximate the irreversible nature of a covalent warhead.

#### Influence of Cellular Protein Concentrations

The cellular context, particularly protein concentrations, plays a crucial role. For both PROTAC types, the model confirms that degradation becomes less efficient with higher initial concentrations of the target protein or with weaker target affinity (**Figure S4A-B**). Encouragingly, the hook effect caused by suboptimal affinity can be mitigated by higher cellular E3 ligase concentrations. In all simulated cases, increasing E3 ligase levels relieved the hook effect, as a larger pool of E3 is available to drive productive turnover (**Figure 7D-F; Figure S3A**). This suggests that the choice of E3 ligase and its expression level in the target cell type are critical variables.

Finally, the model suggests that if the ubiquitinated target can be degraded while still bound in the ternary complex, or if the ubiquitinated ternary complex (TubPL, TubP*L) is inherently destabilized (indicated by a high “destabilization” parameter), then a higher rate of ternary complex formation (cooperativity, α) would be advantageous (**Figure S3B**). All our simulations are based on a high “destabilization” parameter, as this coincides with experimental observations that high cooperativity is generally beneficial.

## Discussion

In this study, we developed PSIRC3, a potent covalent BTK PROTAC that serves as a mechanistically unambiguous tool to resolve the long-standing debate surrounding covalency in TPD. By using a warhead devoid of confounding non-covalent affinity, we have established an unequivocal link between covalent bond formation at Cys481 and efficient BTK degradation.

Our findings directly challenge the dogma that irreversible target binding is incompatible with the PROTAC mechanism. While PSIRC3 is *stoichiometric* with respect to the target (one PROTAC molecule per target molecule), its remarkable potency suggests that *catalytic turnover of the PROTAC itself* is not a prerequisite for high efficacy. The mechanism appears to follow a dynamic, two-stage process. Initially, free PSIRC3 may rapidly and reversibly engage CRBN, leading to non-productive binary complexes and a transient hook effect at high concentrations. Concurrently, PSIRC3 irreversibly modifies BTK, generating a stable BTK–PSIRC3 substrate pool. As free PSIRC3 dissociates from CRBN (or as new CRBN is available), the liberated ligase can re-engage this covalent substrate, forming a productive ternary complex and driving degradation. This “E3-catalysis” model, where the PROTAC is stoichiometric but the E3 ligase is catalytic, explains both the transient hook effect and its resolution over time.

Importantly, insights from our computational modeling further clarify the design principles. For covalent PROTACs, E3 ligase affinity is not a simple “tighter-is-better” relationship—excessively strong binding induces a severe hook effect. In contrast, higher target affinity is generally beneficial. Interestingly, non-covalent PROTACs display an opposite tolerance: weak warheads can coexist with strong E3 ligase binders, whereas strong non-covalent warheads mimic covalent behavior. Across both classes, higher E3 cellular abundance can mitigate hook effects. These findings emphasize that the success of covalent PROTACs rests on achieving a delicate balance between target affinity, E3 affinity, and ternary complex dynamics.

Our comparative analysis demonstrates that the strength of ternary complex formation is the main predictor of covalent PROTAC potency. In BTK-PROTAC-CRBN complex formation assays, PSIRC3 consistently produced a robust, concentration-dependent ternary signal, correlating with its superior degradation efficacy. These findings indicate that optimizing ternary complex cooperativity and productivity—not just cell permeability or target engagement—is crucial for effective covalent protein degradation. The data support prioritizing ternary complex formation in the rational design of potent covalent PROTACs.

This work provides crucial mechanistic clarity that was missing from previous studies. We demonstrate that covalent PROTACs are a powerful and viable strategy, particularly for expanding the druggable proteome. For proteins lacking deep binding pockets or resistant to high-affinity reversible binders, covalent warheads can stabilize weak interactions and convert non-functional binders into potent degraders. This principle mirrors the success of covalent inhibitors for targets like KRAS(G12C) and can now be extended into the degrader space. Advances in chemoproteomics are accelerating the discovery of new covalent ligands beyond cysteine, targeting residues such as lysine, tyrosine, serine, and histidine. By integrating these novel warheads with rational degrader design, it is now possible to pursue the degradation of a much broader array of disease-relevant proteins.

The principle that a stoichiometric, proximity-inducing molecule can be highly effective is not limited to protein degradation. This concept is mirrored by the recent emergence of other non-catalytic modalities. For example, Regulated Induced Proximity Targeting Chimeras (RIPTACs) are heterobifunctional molecules that induce a stable ternary complex between a target protein and a pan-expressed essential protein (e.g., BRD4), which stoichiometrically abrogates the essential protein’s function and leads to selective cell death^46,47^. Similarly, the “CellTrap” mechanism uses a bifunctional molecule to leverage a highly abundant “presenter” protein (like FKBP12) to enrich the molecule intracellularly, thereby potentiating its inhibitory effect on a target like BRD4 ^48^. Perhaps the most striking example is the design of molecules that remodel the surface of Cyclophilin A (CYPA) to create a neomorphic interface, enabling high-affinity, selective binding to the active state of “undruggable” oncogenes like KRAS G12C. This strategy, which results in a stable, inhibitory CYPA:drug:KRAS tricomplex, has shown tumor regression in preclinical models and is now in clinical trials ^49^. All of these strategies, like our covalent PROTAC, are non-catalytic and rely on the formation of a stable, cooperative ternary complex rather than on small-molecule turnover. This growing body of evidence strongly reinforces our central finding: that driving stable, cooperative ternary complexes is a powerful and viable therapeutic strategy in its own right, independent of a catalytic mechanism.

In conclusion, PSIRC3 provides both mechanistic clarity and pharmacological potency, demonstrating that covalent PROTACs are not only viable but also highly effective. By revealing the kinetic and structural principles that govern their function, this work lays the foundation for the rational design of next-generation covalent degraders, significantly expanding the scope of targeted protein degradation in drug discovery.

## Data Availability

The mass spectrometry raw files for DIA proteomics have been deposited in the MassIVE dataset under accession number MSV000099557.

[https://massive.ucsd.edu/ProteoSAFe/dataset.jsp?accession=MSV000099557].

## Acknowledgments

This research was supported in part by the National Institutes of Health (R01-CA250503 to J.W. and J.A.W.), the Cancer Prevention & Research Institute of Texas (CPRIT, RP220480 to J.W.), Michael E. DeBakey, M.D., Professor in Pharmacology (to J.W.), Center for NextGen Therapeutics seed funding (to J.W.), and American Foundation of Pharmaceutical Education Pre-Doctoral Fellowship (to A.D.M.). J.A.W. is a Clinical Scholar of Blood Cancer United.

## Author Contributions

R.C., H.L., X.Q., and J.W. designed the study. R.C, H.L., X.Y., S.M., A.D.M, and X.Q. conducted the experiments. R.C, H.L. analyzed the data. R.C, H.L. and J.W. drafted the manuscript. All authors read and approved the final manuscript.

## Conflict of interests

The authors declare the following competing financial interest(s): J.W. is a co-founder of Chemical Biology Probes, LLC. and serves as a consultant for CoRegen Inc. J.W. and X.Y. are co-founders of Fortitude Biomedicines, Inc. and hold equity interest in this company. The remaining authors declare no competing interests. J.A.W. consults for AstraZeneca, AbbVie, BeOne, Genentech, Johnson & Johnson, Loxo@Lilly, Merck, and Newave. The remaining authors declare no competing interests.

---

**Figure S1.**
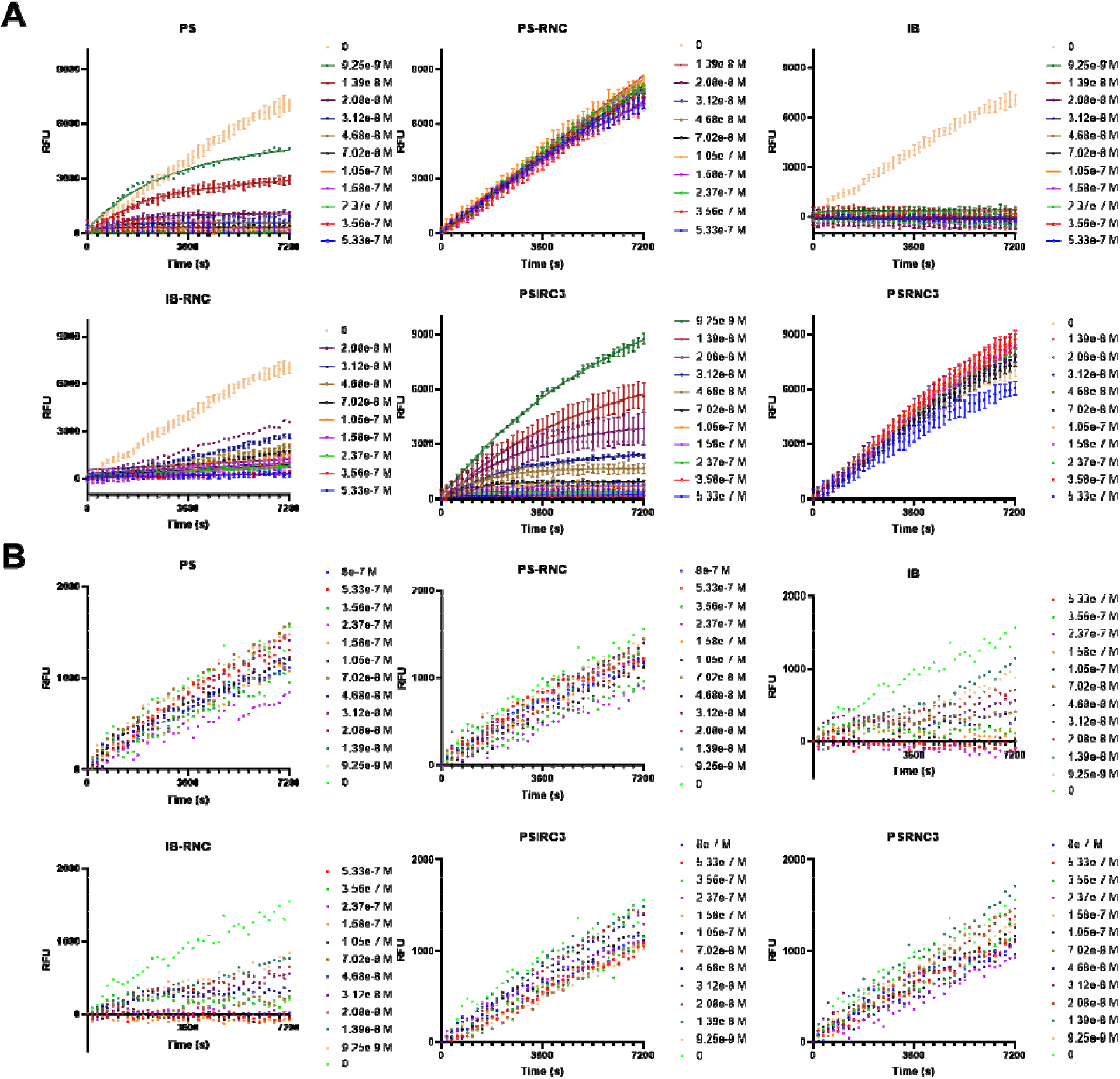
BTK Kinase Activity Assay Confirmed Poseltinib Series are Covalency-driven. The biochemical WT-BTK and C481S-BTK inhibition (BTK Inhibition IC50) was measured using the kinase assay kit from AssayQuant Technologies Inc. (A) WT-BTK, (B) C481S-BTK.

**Figure S2.**
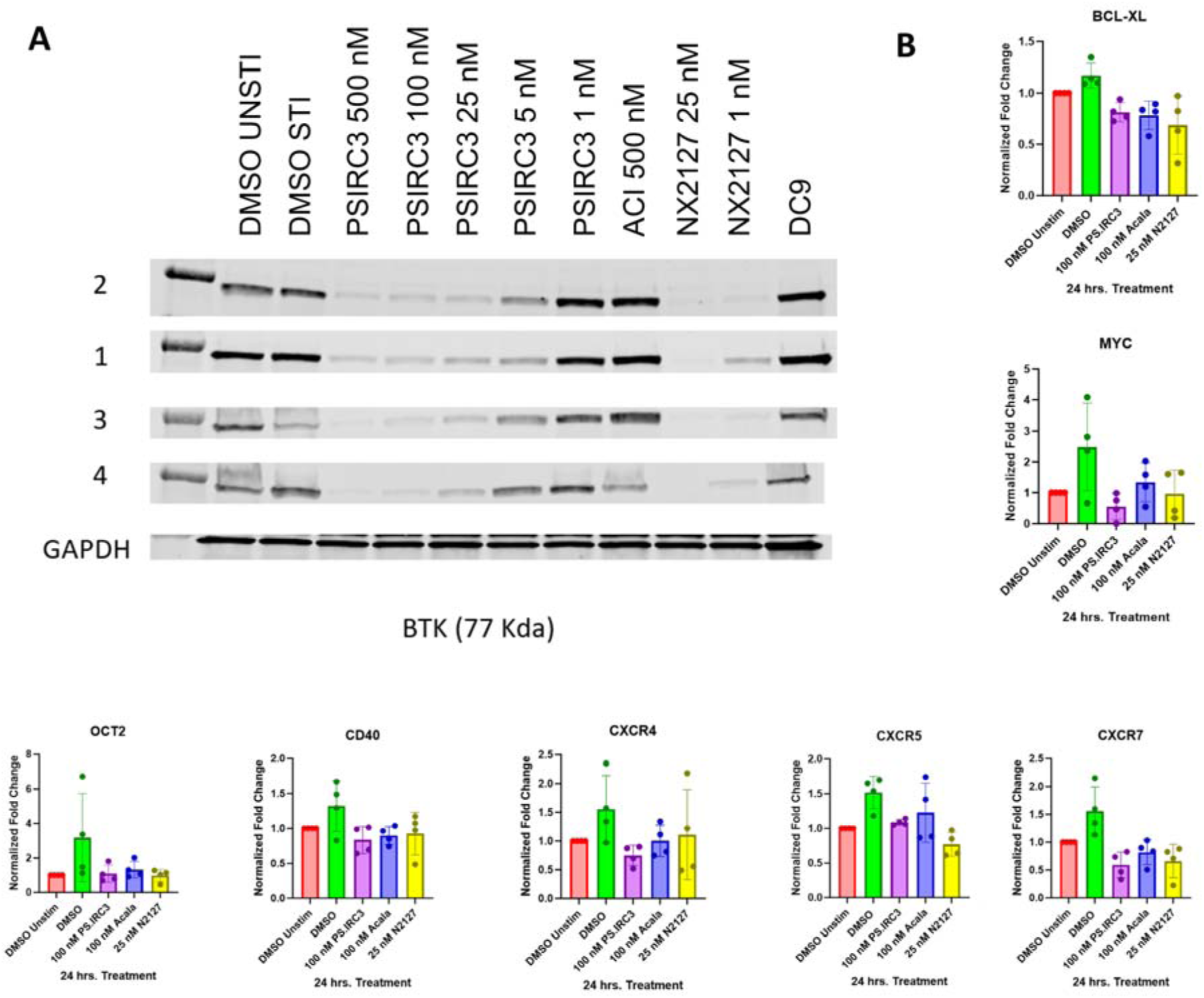
Characterization of BTK Degraders in CLL patient samples. **(A).** PSIRC3 induced degradation of BTK BCR in CLL patient B cells (n=4) analyzed by western blot. **(B).** qRT-PCR analysis of BTK downstream signaling genes of CLL patient B cells (n=4) treated with PSIRC3 for 24 hours with Acalabrutinib and NX-2127 as positive controls.

**Figure S3.**
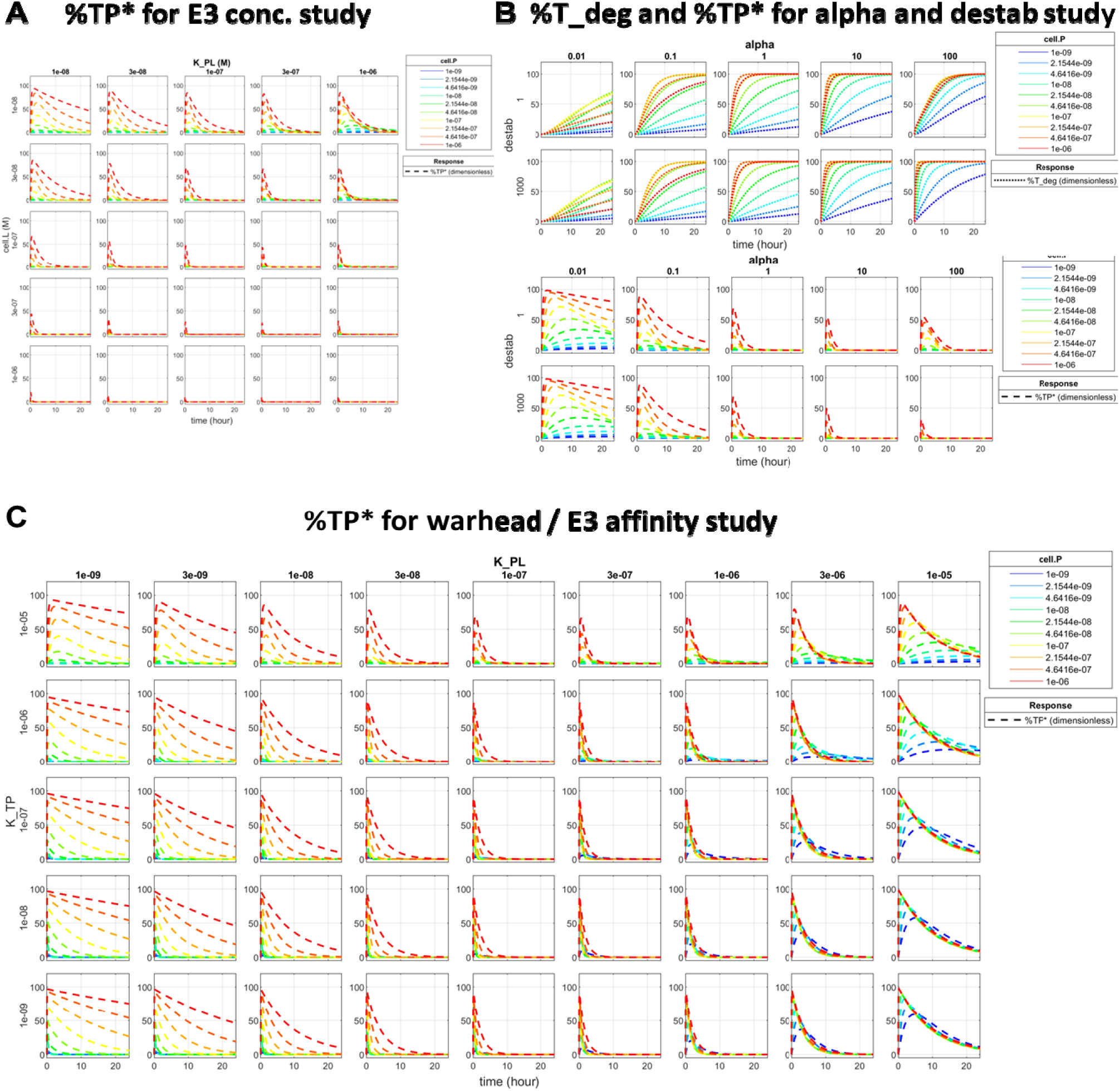
SimBio Modeling of Covalent PROTAC Degradation Event. **(A)** The percentage of TP* (%TP*) associated with the experiment in Figure 5D. **(B)** The time-course simulation of %T_deg and %TP* with various alpha and destab values. **(C)** The percentage of TP* (%TP*) associated with the experiment in Figure 5C.

**Figure S4.**
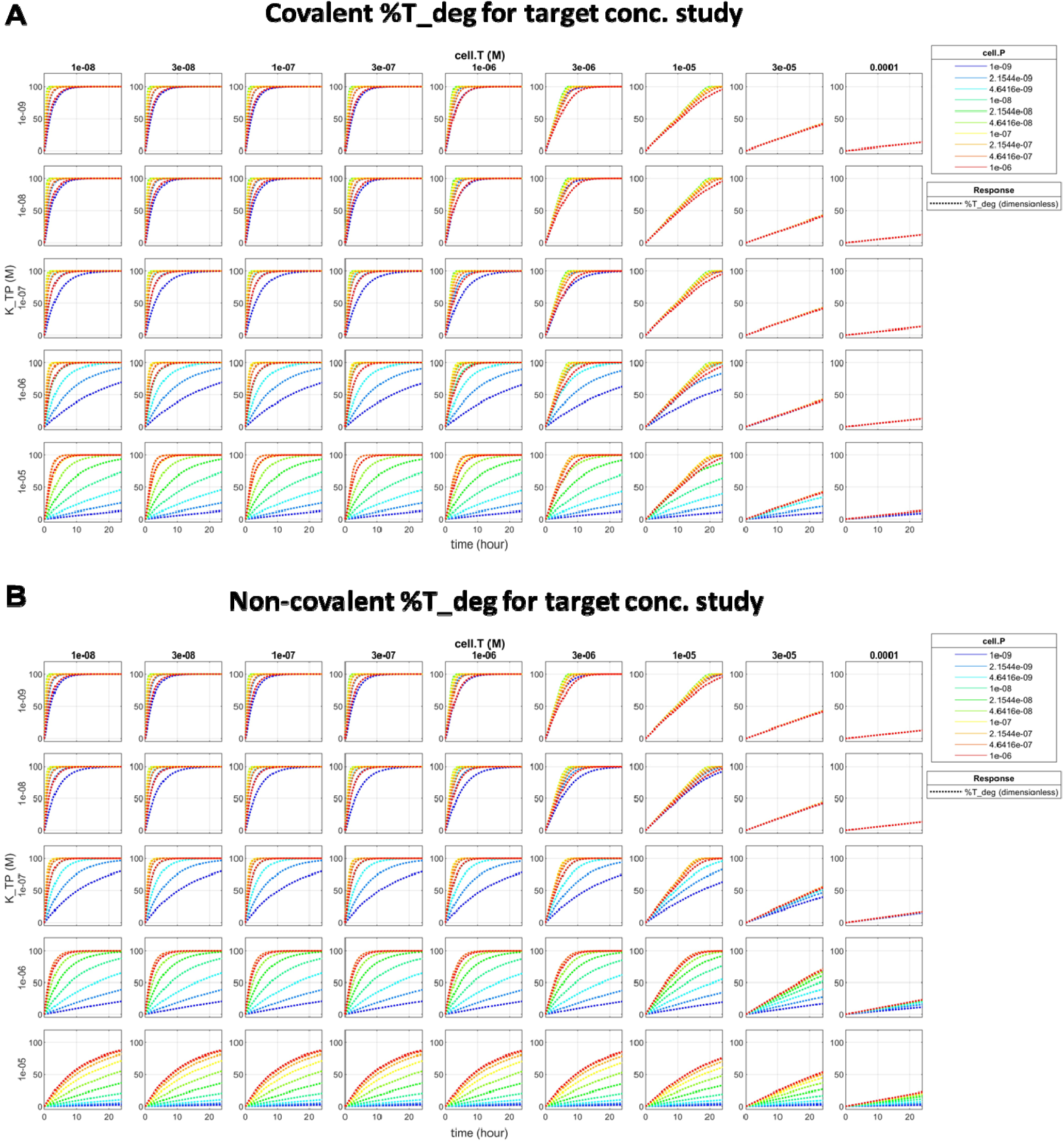
SimBio Modeling of Covalent or Non-covalent PROTAC Degradation Event. The percentage of degraded target protein (%T_deg) with various concentrations of target protein (T) in **(A)** covalent or **(B)** non-covalent PROTAC model.

**Figure S5.**
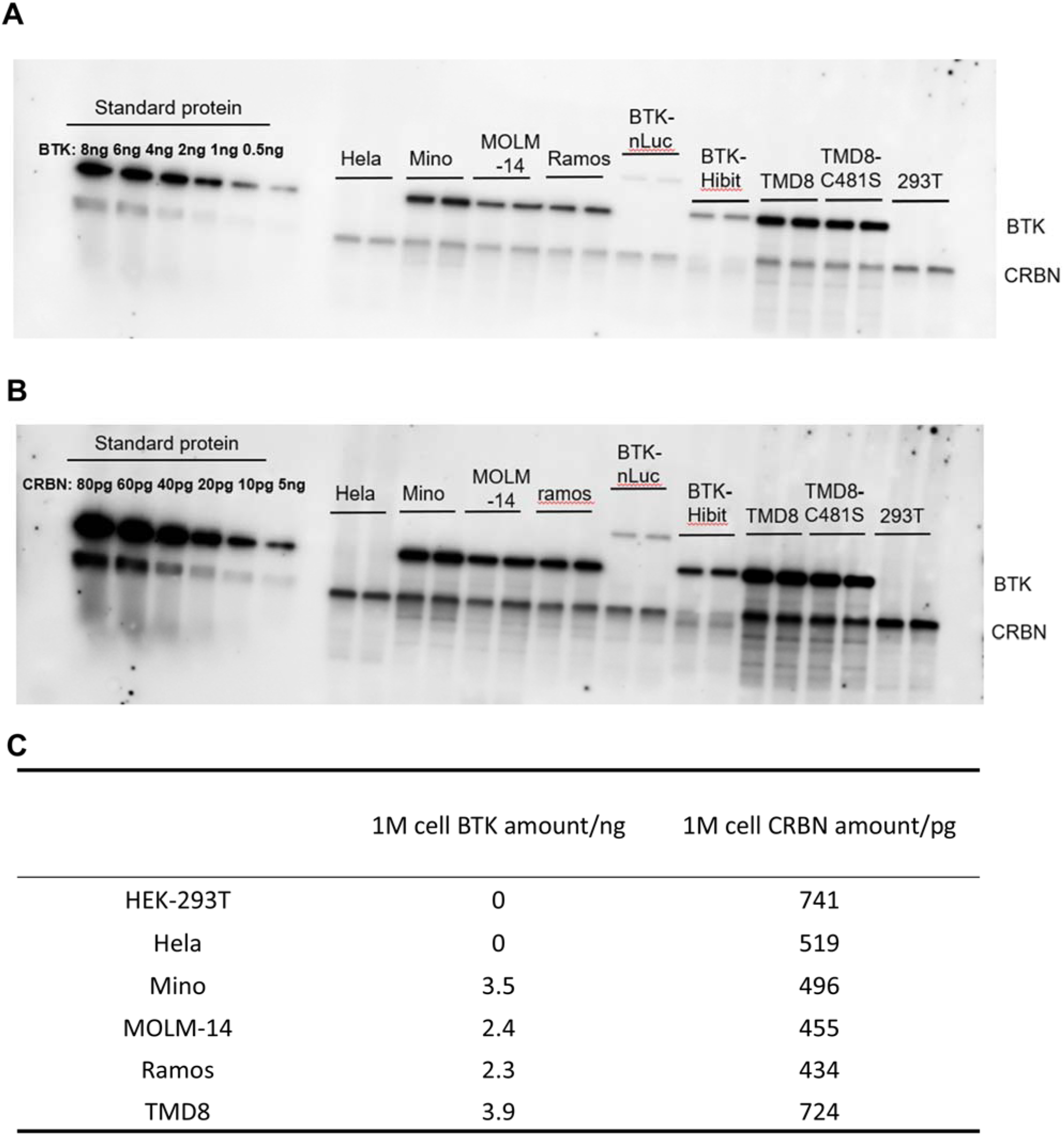
Quantification of BTK and CRBN Protein Levels Across Cell Lines. Quantitative Western blot analysis to determine the endogenous concentrations of BTK and CRBN. **(A)** BTK quantification using a 30s exposure against a standard protein curve. **(B)** CRBN quantification using a 2min exposure. **(C)** Table summarizing the absolute amounts of BTK (ng) and CRBN (pg) per $10^6$ cells across various leukemia and solid tumor cell lines.

